# DADA2: High resolution sample inference from amplicon data

**DOI:** 10.1101/024034

**Authors:** Benjamin J Callahan, Paul J McMurdie, Michael J Rosen, Andrew W Han, Amy Jo Johnson, Susan P Holmes

## Abstract

Microbial communities are commonly characterized by amplifying and sequencing target genes, but errors limit the precision of amplicon sequencing. We present DADA2, a software package that models and corrects amplicon errors. DADA2 identified more real variants than other methods in Illumina-sequenced mock communities, some differing by a single nucleotide, while outputting fewer spurious sequences. DADA2 analysis of vaginal samples revealed a diversity of *Lactobacillus crispatus* strains undetected by OTU methods.

---

The importance of microbial communities to human and environmental health has motivated methods for their efficient characterization. The most common, and cost-effective, method is the amplification and sequencing of targeted genetic elements. *Amplicon sequencing* of taxonomic marker genes such as 16S rRNA [1], the ITS region [2] or 18S rRNA [3] provides a census of a community. Functional diversity can be probed by targeting functional genes [4].

Disentangling errors from biological variation in amplicon sequencing data presents unique challenges, which has prompted the development of amplicon-specific error-correction methods [5, 6, 7, 8]. Most of these methods were designed for pyrosequenced amplicons, and cannot be applied to Illumina sequencing.

Currently, errors in Illumina-sequenced amplicon data are most often addressed by filtering low quality reads and constructing Operational Taxonomic Units (OTUs): clusters of sequences that differ by less than a fixed dissimilarity threshold (typically 3%) within which sequence variation is ignored [9, 10, 11]. Lumping similar sequences together reduces the rate at which errors are misinterpreted as biological variation, but OTUs under-utilize the quality of modern sequencing by precluding the possibility of resolving *fine-scale* (or *strain-level*) variation [7, 12, 13, 14, 15]. Recent studies have shown that fine-scale variation can be informative about ecological niches [12, 13], temporal dynamics [15], and population structure [4]. Fine-scale variation differentiates pathogenic from commensal strains in some cases [16, 17], and can contain clinically relevant information for more complex microbiome-associated diseases [18, 19, 20].

DADA - the Divisive Amplicon Denoising Algorithm - was introduced to correct pyrosequenced amplicon errors without constructing OTUs [7]. DADA was shown to identify real variation at the finest scales in 454-sequenced amplicon data while outputting few false positives [7, 4].

Here we present DADA2, an extension and reimplementation of DADA adapted for use with Illumina sequencing and available as an open-source R package available at https://github.com/benjjneb/dada2. DADA2 implements a new model of Illumina-sequenced amplicon errors that incorporates quality information. Banded alignments and a kmer-distance screen improve computational performance. The DADA2 R package provides light-weight tools for other key parts of the amplicon denoising workflow: filtering, dereplication, chimera identification, and merging paired-end reads.

We compared DADA2 to two other algorithms (Methods): UPARSE, an OTU-construction algorithm with the best published false-positive results [11], and MED, an algorithm with the best published resolution of fine-scale variation in Illumina-sequenced amplicon data [14].

We used three test datasets: Balanced, HMP and Extreme (Methods, Table S1). These data were sequenced at a depth of over 500, 000 highly overlapping paired-end Illumina MiSeq 2x250 reads. The Balanced community consisted of 57 bacteria and archaea mixed at nominally equal frequencies [21]. The HMP community consisted of 21 well-separated bacteria mixed at nominally equal frequencies [22]. The Extreme community consisted of 27 bacterial strains from the human gastrointestinal tract mixed at frequencies spanning five orders of magnitude and with 16S sequences separated by as little as one nucleotide (Methods, Table S2). Sequence quality varied: Balanced was higher (Mean Q = 35.9 forward/33.5 reverse), Extreme moderate (33.0/29.3), and HMP lower (32.3/28.7).

We evaluated specificity by BLAST-ing output sequences against the nr/nt database (Methods). Output sequences with an exact match (100% identity, 100% coverage) were classified as a “Match”, those with a best hit containing one mismatch or one gap were classified as “One Off”, everything else was classified as “Other”.

We evaluated sensitivity by matching output sequences to the 16S sequences from each reference strain. Of note, the number of reference *strains* did not typically match the number of reference *sequences*: some reference strains were identical over the 16S region sequenced, while the genomes of others contained multiple 16S variants. Within-genome variation was useful diagnostically: fine-scale variation was present even in mock communities chosen to be well-separated.

We compared the sample sequences output by DADA2 to the representative sequences output by UPARSE (Figure 1). Almost all variants with Hamming separation greater than the OTU radius (3%, dashed line) were identified by both algorithms (black). However, DADA2 also revealed the fine-scale variation: DADA2 identified biological variants that UPARSE did not (blue) within the OTU radius, in both the merged reads (Figure 1) and the forward reads alone (Figures S1-S3). Both algorithms identified low-frequency variants present in as few as two reads.

**Figure 1.**
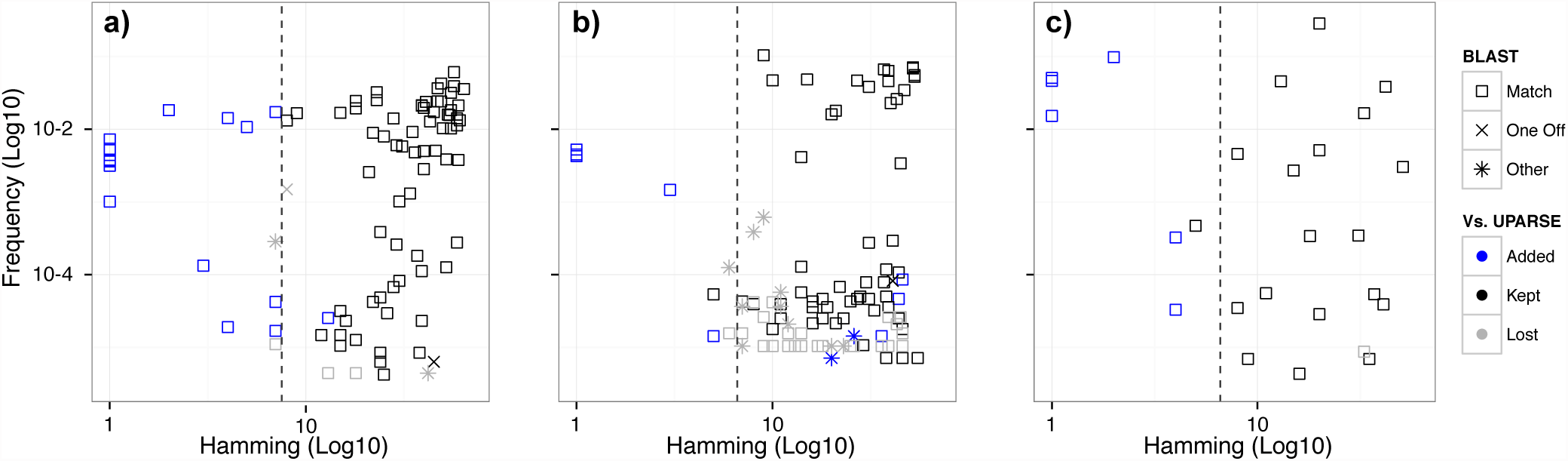
Sample sequence inferred by DADA2 relative to the OTUs constructed by UPARSE. The sequences output by DADA2 are plotted for three Illumina-sequenced amplicon datasets:(a) Balanced, (b) HMP, and (c) Extreme. Frequency is plotted on the y-axis. Hamming distance from each sequence to its closest more-abundant neighbor is plotted on the x-axis. Sample sequences inferred by DADA2 highly overlap with the OTU representative sequences output by UPARSE (black). DADA2 also detects additional biological variation (blue), especially within the OTU radius (dashed line), while outputting fewer spurious sequences (Other).

DADA2 identified more reference sequences and as many or more reference strains than UPARSE in every dataset, whether merging reads or using forward reads alone (Table 1). DADA2 identified every reference strain in the Balanced and HMP datasets; the Extreme reference strains it missed illustrate its limits (SI Note 1).

**Table 1.** The output of DADA2, UPARSE and MED on the test datasets Balanced, HMP and Extreme. Algorithms were applied to the forward reads alone, and the merged forward and reverse reads for each filtered dataset. Output sequences were evaluated by BLAST: Match indicates an exact hit, One Off indicates the top hit has one mismatch, Other indicates larger differences. Mock communities consist of a mixture of reference strains. The number of unique 16S sequences from those strains output by each algorithm and the percentage of the reference strains detected by each algorithm are listed.

|  |  | Reads Denoised | Denoised Sequences |  |  |  | Reference |  |
| --- | --- | --- | --- | --- | --- | --- | --- | --- |
|  |  |  | Total | Match | One Off | Other | Sequences | Strains (%) |
| Balanced | DADA2 (forward) | 99.2% | 94 | 93 | 1 | 0 | 57 | 100.0% |
|  | DADA2 (merged) | 96.2% | 87 | 86 | 1 | 0 | 55 | 96.5% |
|  | UPARSE(forward) | 95.5% | 81 | 77 | 2 | 2 | 53 | 84.2% |
|  | UPARSE (merged) | 91.7% | 76 | 72 | 2 | 2 | 50 | 78.9% |
|  | MED (forward) | 94.4% | 85 | 65 | 20 | 0 | 57 | 100.0% |
|  | MED (merged) | 91.0% | 64 | 63 | 1 | 0 | 54 | 94.7% |
| HMP | DADA2 (forward) | 94.2% | 151 | 135 | 9 | 7 | 23 | 100.0% |
|  | DADA2 (merged) | 92.2% | 67 | 64 | 1 | 2 | 23 | 100.0% |
|  | UPARSE(forward) | 85.4% | 164 | 141 | 11 | 12 | 20 | 100.0% |
|  | UPARSE (merged) | 63.1% | 93 | 81 | 1 | 11 | 20 | 100.0% |
|  | MED (forward) | 81.1% | 88 | 49 | 39 | 0 | 23 | 100.0% |
|  | MED (merged) | 64.9% | 33 | 27 | 5 | 1 | 23 | 100.0% |
| Extreme | DADA2 (forward) | 99.5% | 68 | 61 | 3 | 4 | 25 | 85.2% |
|  | DADA2 (merged) | 97.6% | 26 | 26 | 0 | 0 | 23 | 77.8% |
|  | UPARSE(forward) | 86.4% | 74 | 61 | 0 | 13 | 21 | 77.8% |
|  | UPARSE (merged) | 67.6% | 23 | 22 | 0 | 1 | 18 | 66.7% |
|  | MED (forward) | 89.2% | 95 | 44 | 50 | 1 | 15 | 48.1% |
|  | MED (merged) | 67.9% | 32 | 23 | 7 | 2 | 16 | 51.9% |

The sensitivity of DADA2 to fine-scale variation did not come at the cost of lower specificity. DADA2 output fewer “Other” sequences than UPARSE in every dataset (Table 1). For the Balanced forward reads, DADA2 removed 99.998% of substitution errors remaining after filtering. For the HMP forward reads, DADA2 removed 99.8% of errors after filtering, which compares favorably to the best error-correction rate of 92.3% for that dataset reported in Edgar 2015. DADA2 also discarded fewer reads than UPARSE or MED (Table 1).

DADA2 reported more accurate abundances for some variants than did UPARSE (Figure S7). UPARSE greedily adds reads to the OTU of a more abundant sequence if those reads differ from that sequence by less than the OTU radius (i.e. 3%). When biological variants differ by *∼* 3%, UPARSE splits the reads of the less-abundant variant between the more-abundant variant’s OTU (*<* 3% separation) and a new OTU (*>* 3%). Thus, the abundance reported by UPARSE is too high for the more-abundant variant, and too low for the less-abundant variant. DADA2 does not have this problem because it lacks a hard similarity threshold.

MED was developed to distinguish fine-scale diversity in amplicon data [14]. MED has structural similarities to DADA2: both divide amplicon reads into partitions within which the remaining variation is supposed to be artefactual. MED uses a modified single-site minor-allele-frequency threshold to identify real variation, while imposing a minimum abundance requirement to guard against false positives. As a result, while MED was sensitive to fine-scale variation, it had a high false positive rate and did not detect low frequency variants (Table 1, Figures S1-S6). MED’s specificity was better when analyzing merged reads because most of MED’s false positives derived from repeated single site errors, which are reduced in the overlap region by merging.

DADA2 was slower but of comparable speed to UPARSE, and DADA2 easily processed Illumina-scale samples on a laptop. For the filtered Balanced dataset (*∼* 600k forward reads with singletons) UPARSE ran in 10s, DADA2 in 35s, and MED in 2m30s on a 2013 MacBook Pro.

We applied DADA2 to a collection of 157 Illumina-sequenced samples of the vaginal community collected from 42 pregnant women [23]. The vagina is the least diverse human body habitat [1]. It is often dominated by a single Lactobacillus OTU, and the species of that OTU has been used to characterize the community’s state [24]. *Lactobacillus crispatus* is the most common species, and has been associated with stability and good health [24]. DADA2 revealed that the *L. crispatus* community state is more complex than generally recognized: six distinct *L. crispatus* strains were present in multiple samples and substantial abundance (Figure 2a, Table S3).

**Figure 2.**
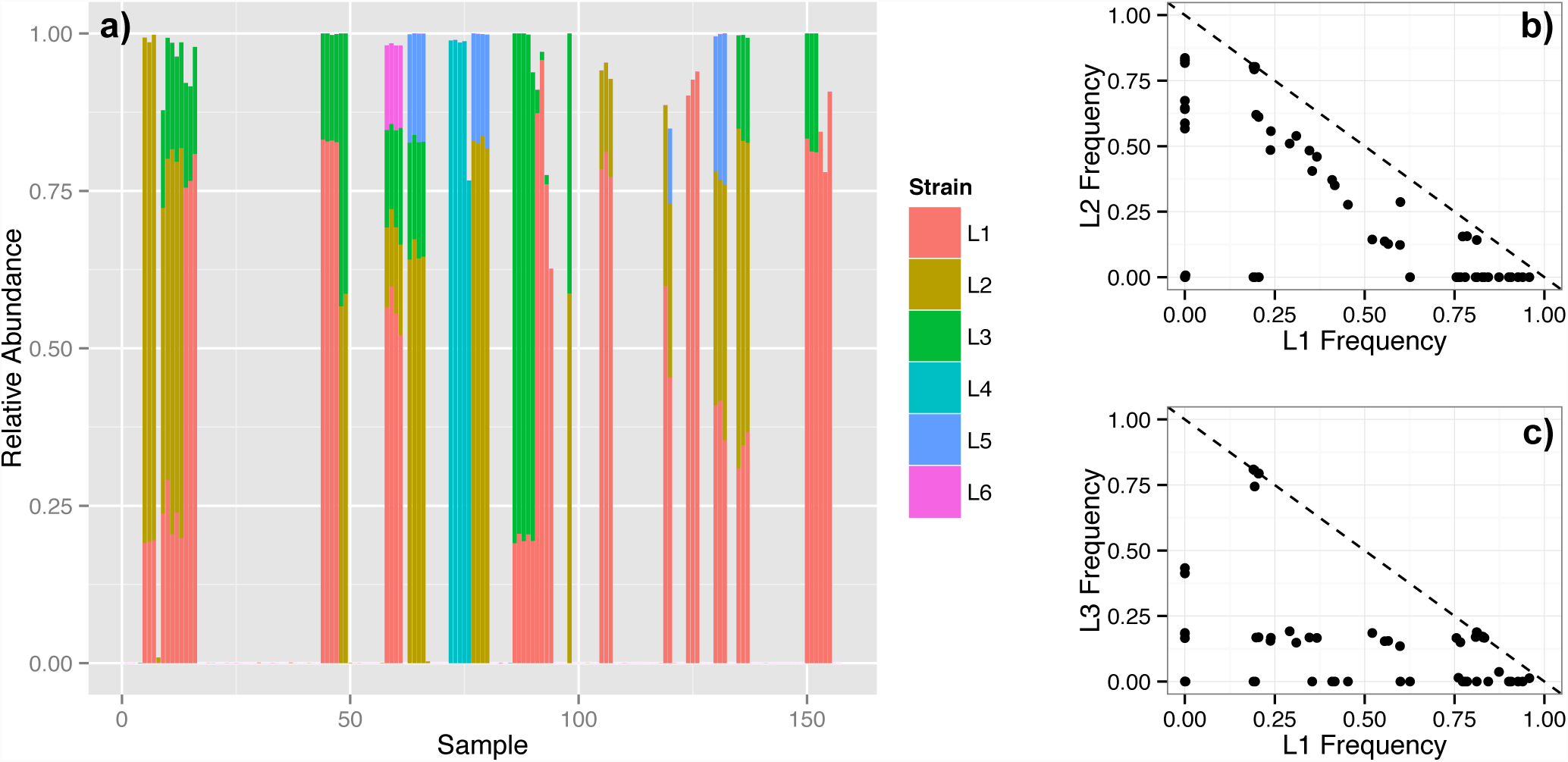
Lactobacillus crispatus strains in the human vaginal community. DADA2 identified six Lactobacillus crispatus strains present in multiple samples and at a significant fraction of all reads (L1: 19.2%, L2: 10.9%, L3: 6.4%, L4: 3.0%, L5: 1.3%, L6: 0.3%). (a) The relative abundance of these strains in each sample. Samples from the same woman are consecutive. The frequency of (b) L1 vs. L2, and (c) L1 vs. L3, for each sample. The dashed line indicates a total frequency of 1.

*L.crispatus* dominated the community when present, but its total abundance was usually split between several strains (Figure 2a). The strain composition of *L. crispatus* dominated communities was stable over time, but substantially differed between women. Of note, the differentiation of *L. crispatus* communities between women is imperceptible to standard OTU analyses that lump together the 16S sequences of these strains that differ by 1-2 nucleotides.

Distinct ecological relationships appear to exist between *L. crispatus* strains. L1 and L2 showed a pattern of mutual exclusion consistent with competition for a common niche (Figure 2b). But L1 and L3 showed a pattern more consistent with a lack of direct competition (Figure 2c): The frequency of L1 was independent of the frequency of L3, which strongly tended towards 20%.

These results show that DADA2 more accurately reconstructs amplicon-sequenced microbial communities. DADA2 better detects fine-scale variation than the current best method for that task, while also outputting fewer incorrect sequences than the most robust OTU method. The precision of DADA2 improves downstream measures of diversity and dissimilarity.

Marker gene sequencing is inherently limited, but the construction of OTUs unnecessarily limits it further. OTUs are not species, and they are not necessitated by amplicon errors. DADA2 makes amplicon sequencing more informative by inferring the composition of amplicon-sequenced microbial communities at the highest resolution.

## Methods

### The Divisive Amplicon Denoising Algorithm

DADA is a divisive partitioning algorithm. All reads begin in a single partition. Reads with the same sequence are grouped into unique sequences with an associated abundance (or dereplicated). The abundance p-value (see next paragraph) is calculated for each unique sequence. If the smallest p-value, after bonferroni correction, falls below the threshold Ω_*A*_, a new partition is formed with that unique sequence as its center. Unique sequences are then allowed to join the partition most likely to have produced them. Division continues until all unique sequences are consistent with being produced as errors from the sequence at the center of their partition, i.e. all abundance p-values are greater than Ω_*A*_. The inferred composition of the sample is the set of central sequences of the final partition and the corresponding number of reads in those partitions (alternatively: each read is denoised by replacing it with the central sequence of its partition).

The abundance p-value quantifies the notion that there are too many reads of a unique sequence *i* for it to be explained by errors in amplicon sequencing. If sequencing errors are independent across reads, the number of reads with sequence *i* that will be produced from sample sequence *j* is poisson distributed with expectation equal to an error rate *λ*_*j→i*_ (see next section) multiplied by the expected reads of sample sequence *j*. Let unique sequence *i* with abundance *a*_*i*_ be in partition *j* containing *n*_*j*_ reads. Then, conditional on I being read at least once, the abundance p-value 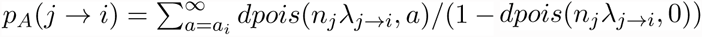. Singletons have an abundance p-value of 1. A low *p*_*A*_ indicates that there are more reads of sequence i than can be explained by errors introduced during the amplification and sequencing of *n*_*j*_ copies of sequence *j*.

### Error rates

DADA2 models errors as occurring independently between sites within a read, and independently between different reads. The probability of a substitution error may depend on the original nucleotide, substituting nucleotide, and associated quality score, e.g. *p*(*A* → *C*, 35). The rate at which read *i* is produced by sequencing sample sequence *j* is then 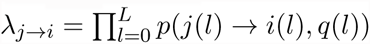.

While DADA2 depends on this parameterized error model, DADA2 does not require prior knowledge of the parameters as it can infer them from the data. After sample inference the substitutions within each partition are tallied and reported to the user by type, allowing the error rates to be estimated. Furthermore, DADA2 implements a self-consist mode which alternates between sample inference and estimating the error rates until the partition and the estimated error rates are jointly consistent.

### DADA2 pipeline

The DADA2 R package implements a complete pipeline to turn paired-end fastq files from the sequencer into merged, denoised, chimera-free, inferred sample sequences. Parts of this pipeline can be substituted with outside methods, but some key elements differ from other approaches.

**Filtering:** fastqFilter() implements filtering of fastq files that largely recapitulates the usearch filterfastq function. fastqPairedFilter() filters paired reads jointly, only outputting reads where both the forward and reverse reads pass the filter.

**Dereplication:** derepFastq() takes an input fastq file and outputs a dereplicated list of unique sequences and their abundances. derepFastq() also outputs position-by-position quality scores for each unique sequence obtained by averaging the positional qualities of the component reads. These averaged scores are used by the error model of the main dada function.

**Denoising:** dada() implements the core denoising algorithm described above.

**Chimeras:** isBimeraDenovo() identifies denoised sequences that are exact bimeras (two-parent chimeras) of more abundant output sequences, or are one-away from an exact bimera of parents that differ from the putative child sequence by at least four substitutions. isShiftDenovo() identifies denoised sequences that are identical to a more abundant sequence up to a shift in starting position.

**Merging:** mergePairs() merges the denoised forward and reverse reads if they exactly overlap. mergePairs() requires that the input forward and reverse reads were in the same order, a feature which is maintained by fastqPairedFilter().

### Test datasets

The Balanced and HMP datasets were downloaded from publicly available sources at http://www.ebi.ac.uk/ena/data/view/PRJEB6244 and http://www.mothur.org/MiSeqDevelopmentData.html. Their construction is described in Schirmer and Kozich respectively.

The Balanced community consists of 57 bacteria and archaea from a broad range of habitats. The 16S sequences of most of these strains were well-separated (*>* 3%) over the region sequenced. However, the 16S sequences of 5 strains were identical to other more abundant strains, while 4 strains had a total of 5 additional distinguishable 16S variants in their genomes that differed by 1 or 2 nucleotides. There were also two strains that were less than 3% different from other more abundant strains.

The HMP community consists entirely of strains which are well-separated (*>* 3%) over the region sequenced. Most of the HMP strains colonize the human body.

The Extreme dataset was generated for this study. The organisms for the Extreme community include human gastrointestinal tract bacterial isolates (Table S2). Strains were chosen to be distinguishable over the 16S region sequenced, but to include a significant amount of fine-scale variation where strains differed by as little as 1 nucleotide from each other.

Extreme strains were grown overnight in liquid broth with the medium recommended from the source culture collection for each respective strain (Table 1). An aliquot of the bacterial culture was used to directly amplify the 16S rRNA gene. One microliter of the bacterial culture was used as template to amplify the V4 region of the 16S rRNA gene using fusion gene primers (515f/806r) that incorporate Illumina adapter sequences and indexing barcodes [25]. The PCR reaction was carried out in a 25 uL mixture containing 1x HotMaster Mix with 2.5 mM Mg2+ (5 PRIME, Gaithersburg, MD), 400 nM forward primer, 400 nM reverse primer, along with the bacterial culture template. The following cycling parameters were used: initial cell lysis and DNA denaturing at 95°C for 10 minutes, followed by 30 cycles of 95°C for 30 seconds, 50°C for 30 seconds, and 72°C for 30 seconds, ending with a final annealing step at 72°C for 10 min. PCR amplicons were cleaned using Agencourt AMPure XP beads (Beckman Coulter, Pasadena, CA) following the manufacturer’s instructions. Cleaned PCR amplicons were analyzed and quantified using an Agilent 2100 Bioanalyzer.

Strains were grouped into two taxonomic groups, Firmicutes and Bacteroidetes. Within each taxonomic group, strains were designated for one of six 10-fold dilution groups (Table S2). PCR amplicons for each strain were first normalized to the same concentration. From there, each amplicon was individually diluted to its respective dilution level and then all amplicons were pooled. The concentration of the pooled library was quantified using the Quant-iT PicoGreen dsDNA Assay kit (Life Technologies, Carlsbad, CA) and analyzed on an Agilent 2100 Bioanalyzer. The pooled library was diluted to 4 nM and then Illumina’s protocol for preparing libraries for sequencing on the MiSeq was followed. The final concentration of the library was diluted to 6 pM with *∼* 20% PhiX spiked in to account for the low base-diversity library. The final pooled library was sequenced on an Illumina MiSeq with a MiSeq Sequencing Reagent Kit v3 to obtain 250 bp paired end reads utilizing custom sequencing primers as described in [25].

### Workflow on test data

A common filtering and trimming was performed before running each algorithm: The DADA2 fastqFilter command was used to remove sequences with Ns or more than two “expected errors” [26], and to trim the first 20 and last 10 (forward) or 10-50 (reverse) bases depending on the quality profile of the data.

The USEARCH command fastq mergepairs with a minimum overlap of 20 bases and maximum differences of 1 was used to merge the filtered forward and reverse reads for further analysis by UPARSE and MED. DADA2 denoised the forward and reverse reads independently, and then merged them with its mergePairs command.

UPARSE and DADA2 require a dereplication step before the main algorithm runs. For UPARSE, the USEARCH command derep fulllength was used for dereplication. For DADA2, the derepFastq command in the R package was used for dereplication. Also, as per the developer’s recommendation, all singletons were removed prior to running UPARSE (but not prior to running DADA2 or MED).

A list of output sequences and associated abundances was obtained for each algorithm. For DADA2 this was the inferred sample sequences, for UPARSE the representative OTU sequences, and for MED the representative sequences of its “nodes”.

Finally, the post-processing tools isBimeraDenovo and isShiftDenovo from the DADA2 R package were used to identify and remove chimeric and shifted sequences from the output sequences of DADA2 and MED. UPARSE does not require this post-processing as it has built-in chimera checking.

Software versions used: USEARCH version 8.0.1623, MED version 2.0, DADA2 version 0.4.3.

### Specificity

Specificity was measured by BLASTing output sequences against the nr/nt database. If the best hit was an exact match covering the full output sequence, it was assigned as a Match. If there was a single mismatch or indel, it was assigned as a One Off. Otherwise it was assigned as Other.

We BLASTed against nr/nt rather than the reference sequences alone because even data sequenced from communities with a putatively known reference composition will contain contaminant sequences. Contaminants are real, albeit unwanted, biological variation, and should be identified when correcting amplicon errors.

While the nr/nt database is imperfect, it is reasonable to expect that Matches are far more likely to be real variants than are Others. Output sequences classified as Other, and output sequences classified as One Off that differed by one substitution from another denoised sequence, were considered as a proxy for false positives.

### Sensitivity

We compiled the 16S sequences (reference sequences) for the intended members of each mock community (reference strains). The presence of each reference strain was confirmed by checking that at least one read of a 16S sequence from each strain was present in the filtered dataset. If no such read existed, that strain was removed from the reference list (two strains were removed from the listed Balanced reference for this reason).

Output sequences were compared to the list of reference sequences. If any output sequence matched any 16S sequences for a given strain, that reference strain was considered to have been identified.

### Analysis of Vaginal Samples

The samples from MacIntyre 2015 were analyzed with the DADA2 pipeline outlined above. First the fastq files were filtered and trimmed in the same manner as the test datasets. Then each sample was dereplicated, and dada() was run with all samples pooled when estimating the error rate parameters. isBimeraDenovo() was used to remove chimeras.

Those taxa that appeared in at least two samples and at least 0.01% of the total reads were taxonomically identified by BLAST. Further analysis focused on the six *L. crispatus* strains identified by this procedure.

## Acknowledgments

We acknowledge funding from NSF DMS-1162538, NIH R01AI112401 and a Microbiome Seed Grant from Stanford-BioX.

### Author Statements

BC and SH designed the research; BC, PM and MR implemented the algorithm; BC performed the analysis; BC, PM, MR and SH wrote the paper; AH and AJ generated the Extreme dataset.

